# A systematic evaluation of deep learning methods for the prediction of drug synergy in cancer

**DOI:** 10.1101/2022.05.16.492054

**Authors:** Delora Baptista, Pedro G. Ferreira, Miguel Rocha

## Abstract

One of the main obstacles to the successful treatment of cancer is the phenomenon of drug resistance. A common strategy to overcome resistance is the use of combination therapies. However, the space of possibilities is huge and efficient search strategies are required. Machine Learning (ML) can be a useful tool for the discovery of novel, clinically relevant anti-cancer drug combinations. In particular, deep learning (DL) has become a popular choice for modeling drug combination effects. Here, we set out to examine the impact of different methodological choices on the performance of multimodal DL-based drug synergy prediction methods, including the use of different input data types, preprocessing steps and model architectures. Focusing on the NCI ALMANAC dataset, we found that feature selection based on prior biological knowledge has a positive impact on performance. Drug features appeared to be more predictive of drug response. Molecular fingerprint-based drug representations performed slightly better than learned representations, and gene expression data of cancer or drug response-specific genes also improved performance. In general, fully connected feature-encoding subnetworks outperformed other architectures, with DL outperforming other ML methods. Using a state-of-the-art interpretability method, we showed that DL models can learn to associate drug and cell line features with drug response in a biologically meaningful way. The strategies explored in this study will help to improve the development of computational methods for the rational design of effective drug combinations for cancer therapy.

**Author summary:** Cancer therapies often fail because tumor cells become resistant to treatment. One way to overcome resistance is by treating patients with a combination of two or more drugs. Some combinations may be more effective than when considering individual drug effects, a phenomenon called drug synergy. Computational drug synergy prediction methods can help to identify new, clinically relevant drug combinations. In this study, we developed several deep learning models for drug synergy prediction. We examined the effect of using different types of deep learning architectures, and different ways of representing drugs and cancer cell lines. We explored the use of biological prior knowledge to select relevant cell line features, and also tested data-driven feature reduction methods. We tested both precomputed drug features and deep learning methods that can directly learn features from raw representations of molecules. We also evaluated whether including genomic features, in addition to gene expression data, improves the predictive performance of the models. Through these experiments, we were able to identify strategies that will help guide the development of new deep learning models for drug synergy prediction in the future.

## Introduction

The phenomenon of drug resistance is one of the greatest challenges in the fight against cancer. Although many tumors initially respond well to a given treatment, the efficacy of single-drug anti-cancer therapies is often diminished due to the existence of tumor drug resistance mechanisms. Resistance-conferring characteristics may already be present in the tumor cells prior to therapy, or they may arise as an adaptive response of the tumor to the treatment itself [1]. One of the main drivers of resistance is intratumoral heterogeneity. Genomic instability in cancer leads to the emergence of subpopulations of cells within a tumor with distinct characteristics and different sensitivity to drugs. Treatment may exert selective pressure on the cells and select subpopulations possessing characteristics that favor drug resistance, leading to future relapse [2].

Combining multiple treatments instead of administering a single drug can help to reduce drug resistance [3]. Drug combinations may circumvent pre-existing resistance mechanisms more easily and prevent the development of acquired resistance mechanisms [4]. In addition, certain combinations may be more effective than would be expected when taking into account the effects of each of the constituent compounds on their own, a phenomenon called drug synergy. Drug synergy increases treatment efficacy without requiring an increase in drug dosage, potentially avoiding an increase in toxicity as well [5]. Therefore, optimal drug combinations will be those that produce synergistic effects.

Novel effective anti-cancer drug combinations can be discovered using high-throughput cell viability assays. In these assays, a large number of candidate drug combinations are screened at different concentrations across a panel of cancer cell lines and the cellular response to the drug is measured. In recent years, several datasets from large-scale drug screening initiatives have been made publicly available [6–8]. The largest of these is the National Cancer Institute (NCI) A Large Matrix of Anti-Neoplastic Agent Combinations (ALMANAC) project [7], which screened a total of 5,232 pairs of FDA-approved drugs against National Cancer Institute 60 Human Cancer Cell Line Screen (NCI-60), a panel of 60 tumor cell lines that have been extensively characterized at the molecular level [9]. The project uncovered several synergistic drug pairs, including two clinically novel combinations that are currently being evaluated in phase I clinical trials [7].

Despite the existence of these high-throughput technologies, screening all conceivable drug combinations is still infeasible, for both practical and financial reasons [8, 10]. Computational methods could greatly reduce the search space, thus minimizing the experimental effort required to find truly effective anti-cancer drug combinations. ML, for example, can be used to learn functional mappings between very high-dimensional input data and a score that reflects drug combination effects. This makes it a powerful approach to develop models that are able to predict drug synergy based on drug combination screening experiments and other relevant data. Several ML models for drug synergy prediction have been described in the literature [8, 11–15]. Many of these studies used tree-based ML methods, such as random forests (RFs) [11, 12, 14] or gradient boosting [12, 13, 15].

One particular subset of ML that has attracted great interest from researchers in this field is deep learning (DL). These are models composed of multiple processing layers [16], giving them the ability to learn complex, non-linear functions. Furthermore, unlike most traditional ML methods, DL approaches typically do not require extensive feature selection before training, since they have the ability to learn higher-order representations directly from raw input data [17]. Since DL models can handle large amounts of high-dimensional and noisy data, they are good candidates for the development of drug synergy prediction models.

Preuer et al. [18] presented DeepSynergy, a feedforward, fully-connected deep neural network that uses chemical features and gene expression data to predict drug synergy. Xia et al. developed a multimodal DL model to predict the growth inhibition of cell lines from the NCI ALMANAC project [19]. This model includes separate feature-encoding subnetworks for each input data type (drug descriptors, gene expression, microRNA and proteomics data) and a cell line growth prediction network. Several other DL-based drug synergy prediction models have since been reported in the literature. Similar to the model proposed in 2018 by Xia et al., many of these more recent models adopt a multimodal architecture [20–23].

Beyond fully connected models [18, 19, 21, 22, 24], other innovative architectures have been proposed. Zhang et al. [25] developed a sparsely-connected deep belief network constrained by biological prior knowledge. The recent architecture of the TranSynergy model [26] includes a transformer [27] component, as well as fully connected layers. A method called REpresentation of Features as Images with NEighborhood Dependencies (REFINED) was developed to transform drug descriptors into images, so that convolutional neural networks (CNNs) could be used to model drug synergy instead of the typical fully connected networks [28]. Another study used graph neural networks (GNNs) for drug-specific subnetworks to learn drug representations directly from the compound structures in an end-to-end manner [23]. Several recent studies have used GNNs trained on graphs containing information on interactions between the drugs in a combination, between drugs and their targets, and/or interactions between genes or proteins in the cell lines [29–32].

Most drug synergy prediction models use drug features or gene expression features or a combination of both. Other models include additional cell line information, such as genetic data (somatic mutations and/or copy number variations (CNVs)) [20, 25] or proteomics data [19, 24]. Drug target-specific features have also been included [22, 25, 26]. Since adding more features increases the complexity of the models, assessing which types of input data are more informative and predictive of drug synergy is essential.

Precomputed molecular descriptors or fingerprints are used as chemical features to represent the drugs, as an alternative to the use of end-to-end DL methods to learn the relevant compound features directly from the compound structures. Given that the screening datasets that are currently available only contain a very limited number of compounds, it is still unclear whether there is any benefit in using learned representations instead of traditional fingerprints and descriptors. A recent study benchmarked several compound representations on a large drug synergy dataset and found that DL-based representations were able to outperform traditional fingerprints [15]. However, the authors also noted that the difference between the top performing DL-based methods and the best fingerprints was not substantial and that other concerns, such as interpretability, may be more important.

Feature reduction is often applied to the cell line omics data, either by using specific gene lists [23, 25, 26], or by employing unsupervised data dimensionality reduction techniques, such as principal component analysis (PCA) [24, 33] or autoencoders [20, 24]. Using predefined gene lists to select features provides greater control over the selection process and might make the models easier to interpret biologically. However, certain approaches, such as limiting the gene features to known drug targets present in the training set, may limit the generalization of the models. Data-driven approaches avoid this problem.

Another advantage of data-driven dimensionality reduction techniques is the capacity to be trained using much larger datasets with data from more cell lines than those used in the screening datasets [24], or even patient data [20]. Nevertheless, a limitation of this approach is the difficulty in interpreting the results.Therefore, evaluating which feature reduction methods are capable of achieving satisfactory performance, as well as simultaneously facilitating model interpretability, is an essential step when designing drug synergy prediction models.

The impact of different methodological aspects on the drug synergy prediction models is still unclear and a systematic evaluation is missing. In this work, we set out to investigate the impact of different methodological variables on the performance of drug synergy prediction DL methods, using the ALMANAC drug combination screening dataset. Namely, we evaluated the impact of different preprocessing steps, types of input data, and DL architectures on the final performance of the methods. Prior biological knowledge was used to select cell line features and to facilitate model interpretation.

Interpretability is an important requirement of biomedical predictive systems. We further explored recent methodologies to determine the importance of features and the interpretability of the prediction mechanisms.

We were able to identify the types of input data that are more predictive of drug response, as well as the feature selection and data representation methods that produce the best results. We also found that combining different models improves performance. Additionally, we demonstrate that the decisions made by the DL models are driven by biologically meaningful features.

## Results

### Testing the impact of different methodological variables

We developed several multimodal DL models (Fig 1) to predict drug combination effects summarized as ComboScores, using the ALMANAC dataset [7]. The ComboScore for a given *< cell line− drugA− drugB >* triplet is the sum of the differences between the expected and observed cell line growth calculated for each dose combination tested in the screen, with expected effects being determined using a modified version of the Bliss independence reference model [34].

**Fig 1.**
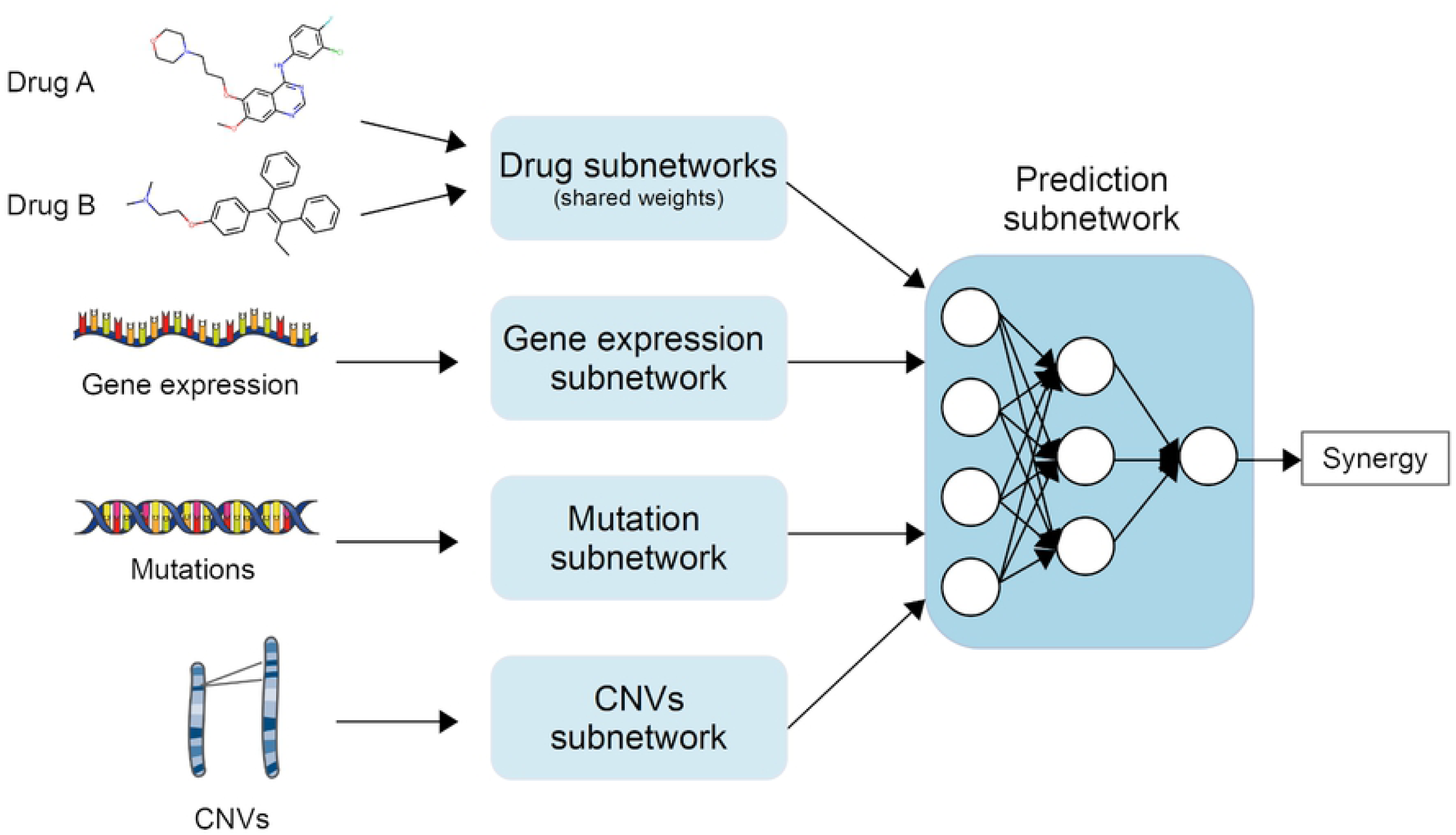
A representation of the general architecture of the multimodal DL models developed in this study. A model is defined as a combination of the learning algorithm itself and the data preparation steps required beforehand.

First, we built several baseline models: a random baseline model that always predicts the average ComboScore value of the training set, and baseline models where one-hot encoded identifiers were used instead of cell line features and/or drug features.

We then built models trained on two different combinations of input data types:

1. *expr + drugA + drugB* models - use of RNA-Seq gene expression data (*expr*) and drug features for both drugs in a combination (drugA, drugB);
2. *expr + mut + cnv + drugA + drugB* - use of expression data (*expr*), mutation data (*mut*), copy number variation features (*cnv*) and drug features (*drugA, drugB*).

We began with type 1 models and tested the impact of the format of the transcriptomics data. We evaluated the influence of the type of architecture used to process the gene expression values (fully connected, 1D CNNs and 2D CNNs) and the influence of the arrangement and order of the genes (*chromosome order* vs. *clustering order*).

We then tested the extension of the gene sets, by using more comprehensive or more selective lists of genes. We also tested if using directly normalized expression values or compact representations that capture the variability and correlation showed any differences in performance. The following feature selection/dimensionality reduction methods were evaluated in this study:

- *expr*_*protein coding*_: the original dataset was filtered so that only protein coding genes were kept, leaving a total of 18,779 genes;
- *expr*_*landmark*_: a list of “landmark genes” as defined by the L1000 project was used to reduce the gene expression dataset to 978 genes that are considered to be representative of the transcriptome as a whole [35];
- *expr*_*DGI*_: Drug-gene interaction (DGI) claims were obtained from The Drug-gene Interaction Database (DGIdb) [36] (version 4.2.0) and then used to build a list of 994 DGI genes for the compounds screened in the ALMANAC project. 976 of these genes were present in the gene expression dataset;
- *expr*_*COSMIC*_: A list of 723 cancer driver genes was obtained from the Catalogue of Somatic Mutations in Cancer (COSMIC) [37] (version 94) Cancer Gene Census [38]. After filtering, the gene expression dataset contained 713 genes;
- *expr*_*NCG*_: A different cancer-specific gene list containing 2,372 genes was obtained from the Network of Cancer Genes (NCG) (version 6.0) [39]. It includes genes identified in the COSMIC Cancer Gene Census, as well as other genes that have been implicated in cancer. After filtering, 2,362 genes remained in the expression dataset;
- Combined gene lists: *expr*_*DGI + landmark*_ (1,815 features) and *expr*_*DGI + NCG*_ (3,037 features);
- *expr*_*UMAP*_: Uniform Manifold Approximation and Projection (UMAP) [40] was used to capture the non-linear structure in the high-dimensional gene expression data, projecting it into a low-dimensional representation (50 components);
- *expr*_*WGCNA*_: Weighted Gene Co-expression Network Analysis (WGCNA) [41] was used to find network modules that capture the correlated expression of a set of genes. Module eigengenes summarize the expression of genes in each module and can be used to reduce the dimensionality of the gene expression data. The reduced expressed dataset had a total of 136 module eigengenes.

Next, we kept the expression-encoding subnetwork fixed and modified the drug-encoding subnetworks to assess the impact of different drug representations:

- *drugs*_*ECFP4*_: 1024-bit Morgan fingerprints with a radius of 2 (equivalent to extended connectivity fingerprint (ECFP)4 fingerprints [42]) fed into fully connected drug subnetworks;
- *drugs*_*LayeredFP*_: 1024-bit layered fingerprints fed into fully connected drug subnetworks;
- *drugs*_*TextCNN*_: Simplified Molecular-Input Line-Entry System (SMILES) strings were tokenized and one-hot encoded, and then fed into TextCNN [43, 44] subnetworks;
- *drugs*_*GCN*_: using graph convolutional networks (GCNs) [45] for the drug subnetworks;
- *drugs*_*GAT*_: using graph attention networks (GATs) [46] for the drug subnetworks;
- *drugs*_*MTE*_: 512-dimensional Molecular Transformer Embeddings (MTEs) [47], fed into fully connected drug subnetworks.

Afterwards, we examined if additional data on mutations and CNVs would lead to improved predictive performance (type 2 models). We tested two different models: one trained on mutation data summarized at the gene level and CNVs, filtered to only include DGI genes (*expr*_*DGI*_ *+ mut*_*DGI, gene-level*_ *+ cnv*_*DGI*_ *+ drugs*_*ECFP4*_ model); a second model trained using mutations summarized at the pathway level and CNVs (*expr*_*DGI*_ *+ mut*_*pathway-level*_ *+ cnv*_*DGI*_ *+ drugs*_*ECFP4*_ model).

We also compared our DL models against several ML algorithms (elastic net, support vector regression (SVR), RF, extreme gradient boosting (XGBoost) and light gradient boosting machine (LGBM)). In total, we developed 24 different DL models and 6 ML models, which are summarized in S1 Fig. The results of these tests will be described in the following subsections.

### Baseline models

The baseline models that were built served as references for subsequent models. All of the models developed in this study performed better than a random baseline model, that always predicts the average ComboScore value of the training set (S1 Table).

To assess the importance of including a certain input data type, we analysed the performance of models trained on one-hot encoded identifiers instead of actual cell line or drug features:

- *cell line*_*one hot*_ *+ drugs*_*one hot*_ - a model trained exclusively with one-hot encoded cell line and drug identifiers, which was used to determine if omics and chemical features include information that is relevant for the prediction of drug synergy, beyond the information contained in drug or cell line identifiers;
- *cell line*_*one hot*_ *+ drugs*_*ECFP4*_ - to determine the impact of removing cell line omics features on the performance of the model;
- *expr*_*DGI*_ *+ drugs*_*one hot*_ - to determine the impact of removing drug features;

Nearly all of the models performed better than the *cell line*_*one hot*_ *+ drugs*_*one hot*_ model (Fig 2). Using one-hot encoded cell line names instead of omics features (*cell line*_*one hot*_ *+ drugs*_*ECFP4*_ model) decreased model performance (Fig 2A). Nevertheless, the scores are comparable to those of some of the models that had been trained on actual gene expression data. When using one-hot encoded drug identifiers instead of chemical features as input (*expr*_*DGI*_ *+ drugs*_*one hot*_ model), model performance dropped even more (Fig 2B). Both the (*cell line*_*one hot*_ *+ drugs*_*ECFP4*_ model and the *expr*_*DGI*_ *+ drugs*_*one hot*_ model performed better than the *cell line*_*one hot*_ *+ drugs*_*one hot*_ model. These results seem to indicate that both drug and gene expression features contribute to the predictive capacity of the model, and that drug features appear to be more predictive of drug synergy than omics features.

**Fig 2.**
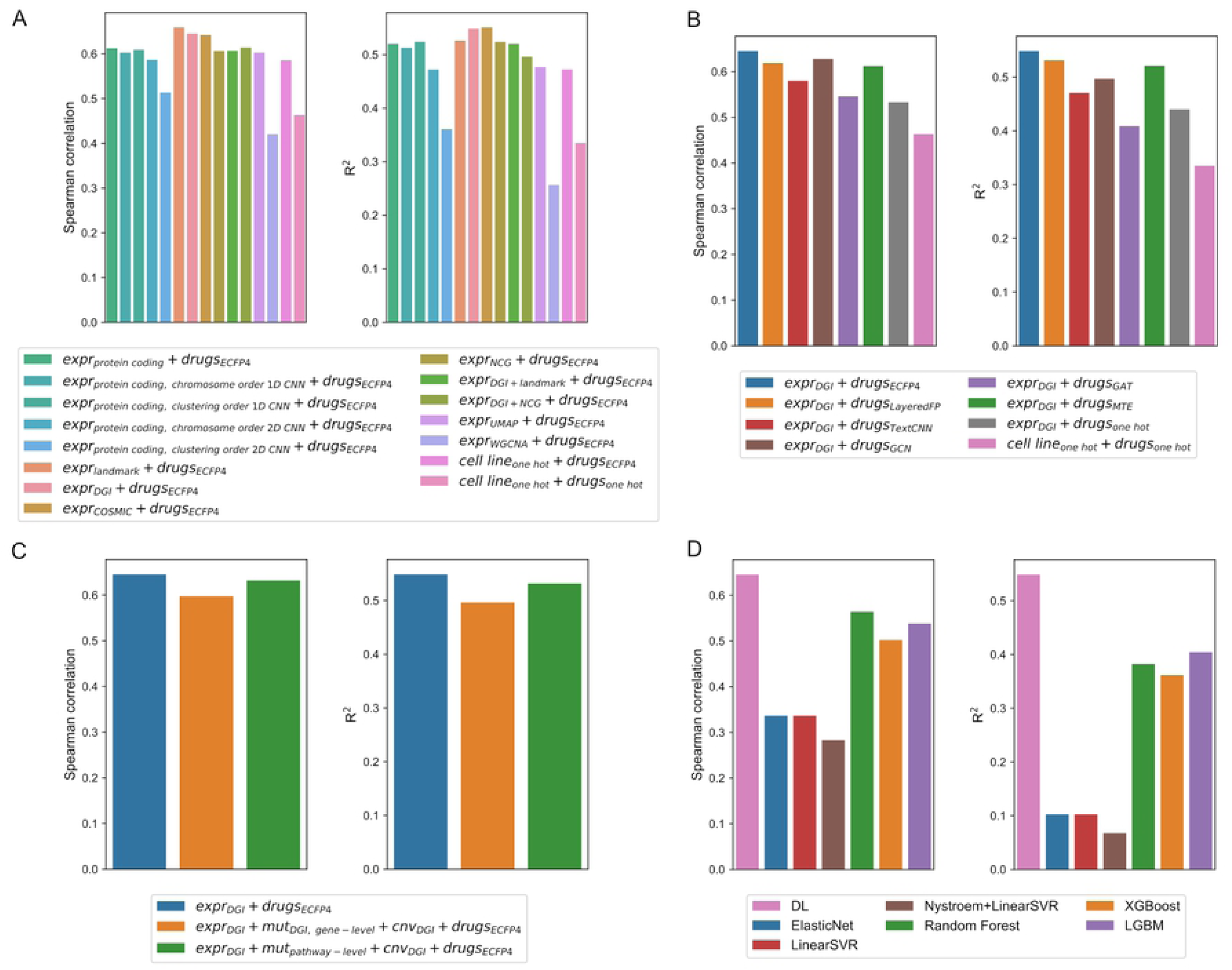
Performance scores (Spearman correlation - left - and *R*^2^ - right) of the different models tested in this study. (A) Performance scores of models with different gene expression feature-encoding subnetworks. (B) Performance scores of models with different drug encoding subnetworks. (C) Performance scores of models trained with and without mutation (*mut*) and copy number variation (*cnv*) data in addition to gene expression and drug features. (D) Performance scores of non-DL models compared with one of the best DL models developed in this study.

### Different gene expression subnetworks

For the most part, models with fully connected gene expression subnetworks trained using the original gene expression values (tabular format) outperformed 1D and 2D convolutional gene expression subnetworks (Fig 2A). The 1D CNN subnetworks were able to achieve performance scores that were similar to some of the lower ranked fully connected gene expression subnetworks, but the 2D CNN subnetworks performed worse. This was independent of the order of genes obtained by clustering or by chromosome position. In light of these results, we only used fully connected layers for the omics subnetworks in subsequent tests.

The best gene expression subnetworks were fully connected subnetworks trained on the log-transformed and min-max scaled Fragments Per Kilobase of transcript per Million mapped reads (FPKM) values, with genes being selected according to predefined gene lists (Fig 2A). The top three models were trained on expression data limited to landmark genes, COSMIC cancer genes or genes with known or potential interactions with drugs screened in the ALMANAC project (*expr*_*DGI*_). Working with the full set of over 18,000 protein coding genes present in this dataset was not shown to be particularly advantageous. Most of the feature selection methods that used smaller gene lists produced models with similar or higher performance scores.

Using dimensionality reduction led to worse performance (Fig 2A). UMAP embeddings were able to perform similarly to some of the gene list selection methods, while using WGCNA as a dimensionality reduction method resulted in poorer performance, worse than the baseline models trained on one-hot encoded cell line names.

In the following sections, we will only focus on models using *expr*_*DGI*_ as the feature-encoding subnetwork for gene expression data. This gene selection method ranked second in terms of scoring metrics (coefficient of determination (R^2^) and Spearman correlation), and might also provide better interpretability.

### Drug encoding networks

The model trained on ECFP4 fingerprints outperformed all of the other subnetworks in terms of all scoring metrics (Fig 2B). Another fingerprint-based drug encoding scheme, RDKit layered fingerprints (LayeredFP), achieved the second-best R^2^ score, while the GCN subnetwork achieved the second-best Spearman correlation score. In general, most of the drug subnetworks achieved similar performance scores, minus a few exceptions (such as the GAT subnetwork).

### Additional cell line features

Including mutation and CNV data slightly decreased model performance (Fig 2C). The model trained on pathway-level mutation features performed slightly better than the model trained using gene-level mutation data. This may be due to the fact that more genes were taken into consideration when summarizing the mutation data at the pathway level, which would be an indication that relevant genetic information is being lost when only DGI genes are considered. The mutation dataset summarized at the pathway-level is also less sparse (i.e. a lower percentage of entries are zero) than the gene-level dataset.

### Comparison with other machine learning models

To determine if there is any advantage in using DL instead of other ML algorithms to model drug combination effects, we compared our DL model trained on gene expression features of DGI genes and ECFP4 fingerprints (*expr*_*DGI*_ *+ drugs*_*ECFP4*_ model) against 5 different ML models trained on the same features (Fig 2D). We tested elastic net, linear SVR and linear SVR preceded by a radial basis function (RBF) kernel approximation method, RF, XGBoost and LGBM models. The DL model outperformed all of the ML models. The best non-DL models were LGBM (in terms of R^2^) and RF (in terms of Spearman correlation). The performance of all of the tree-based models (RF, XGBoost and LGBM) is on par with some of the lower ranking DL models described in previous sections. The elastic net and SVR models did not perform well, with worse performance than the baseline DL model trained on one-hot encoded identifiers.

### Heterogeneous ensemble

We also created a simple heterogeneous ensemble, to determine if combining different DL approaches, as well as other ML models, could lead to better results. The ensemble included the top 10 DL models developed in this study and the RF, XGBoost and LGBM models. To obtain the ensemble prediction, the predictions from each of the individual models were simply averaged.

The heterogeneous ensemble outperformed the best individual DL models for both scoring metrics (R^2^=0.584 and Spearman correlation=0.672 vs. R^2^=0.549 and Spearman correlation=0.645 for the *expr* _*DGI*_ *+ drugs*_*ECFP4*_ model, for example). These results show that combining different DL architectures and different learning algorithms can improve the generalizability of drug synergy prediction models.

### Feature importance

One of the main disadvantages of using DL models to predict drug response is that they are difficult to interpret. The SHapley Additive exPlanations (SHAP) [48] interpretability framework was used to determine which features were the most important for the *expr*_*DGI*_ *+ drugs*_*ECFP4*_ model. SHAP values reflect the contribution of each feature to a prediction.

Fig 3 shows the top 20 most important features. Important features are those that have greater absolute SHAP values. Each point in the figure represents an instance in the test set. The points are distributed along the X-axis according to the SHAP values. The colors represent the feature values. This reveals how the value of a feature influences the model output for a given sample. For example, when ECFP4 bit 250 is “ON” (value=1), the impact on the model output is usually positive, pushing the predicted ComboScore higher.

**Fig 3.**
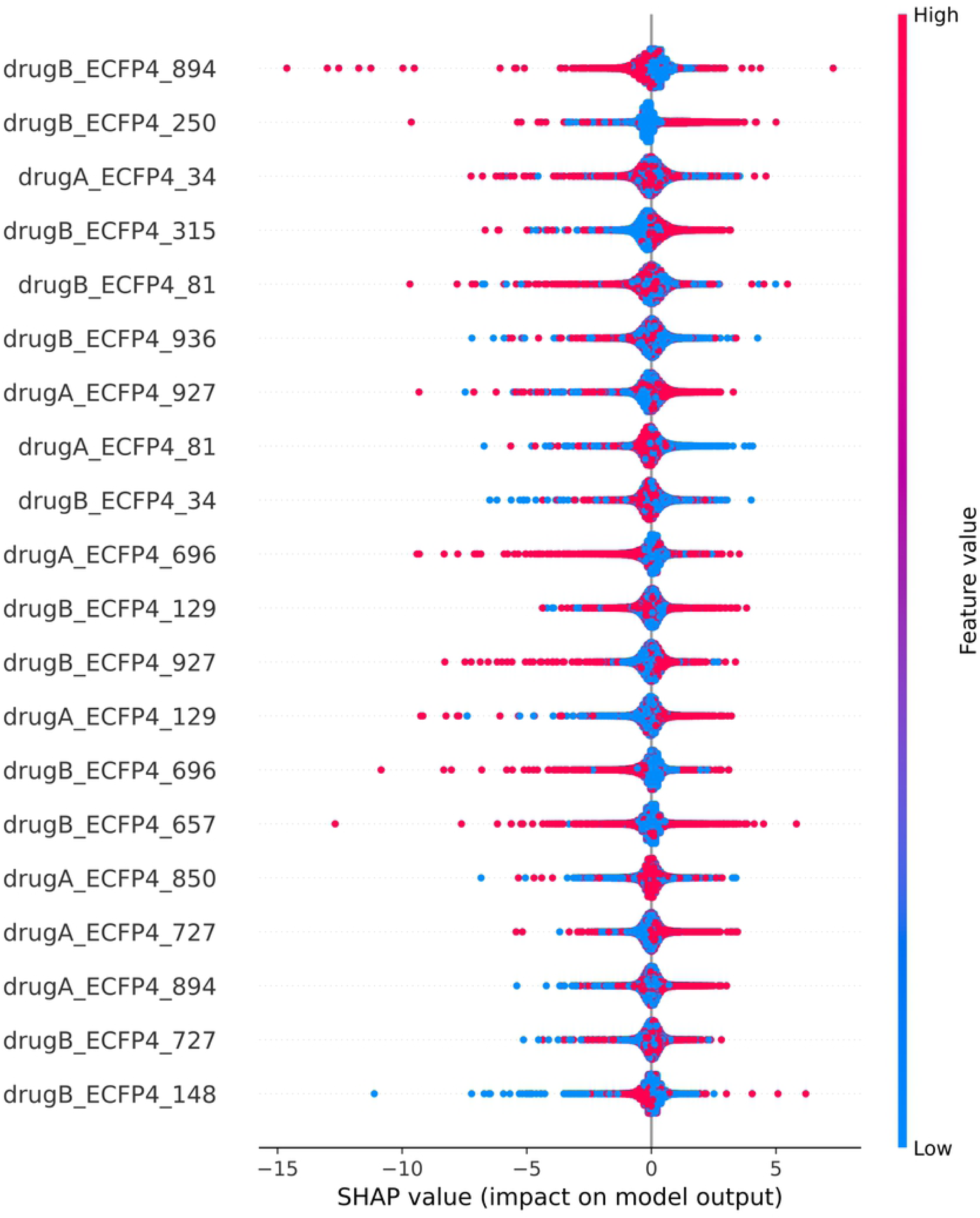
Top 20 most important features, ranked by mean absolute SHAP values.

All of the top 20 features for this DL model are drug features. A similar result is obtained for the RF model (S2 Table), suggesting that, for the ALMANAC dataset, drug features are the most relevant for drug response prediction.

SHAP can also be used to provide explanations for specific examples. We analyzed a specific example (11835) that involved the SR cell line, derived from an anaplastic large cell lymphoma [49]. The drugs screened in this experiment were Topotecan hydrochloride, which interacts with the topoisomerase I-DNA complex at the site where the DNA cleavage occurs [50], and Gefitinib, which inhibits the epidermal growth factor receptor (EGFR) tyrosine kinase domain [51]. This experiment was selected for further analysis since this combination was found to have greater-than-additive effects in the SR cell line in the ALMANAC screen (ComboScore=108.00), and because the ComboScore predicted by our model (ComboScore=108.06) was close to the true value. Fig 4A displays the explanation of the prediction for this case. Each row shows the contribution of each feature to improve the prediction score. For this specific example, all of the top 20 features were, once again, ECFP4 fingerprint bits from drugA and drugB.

**Fig 4.**
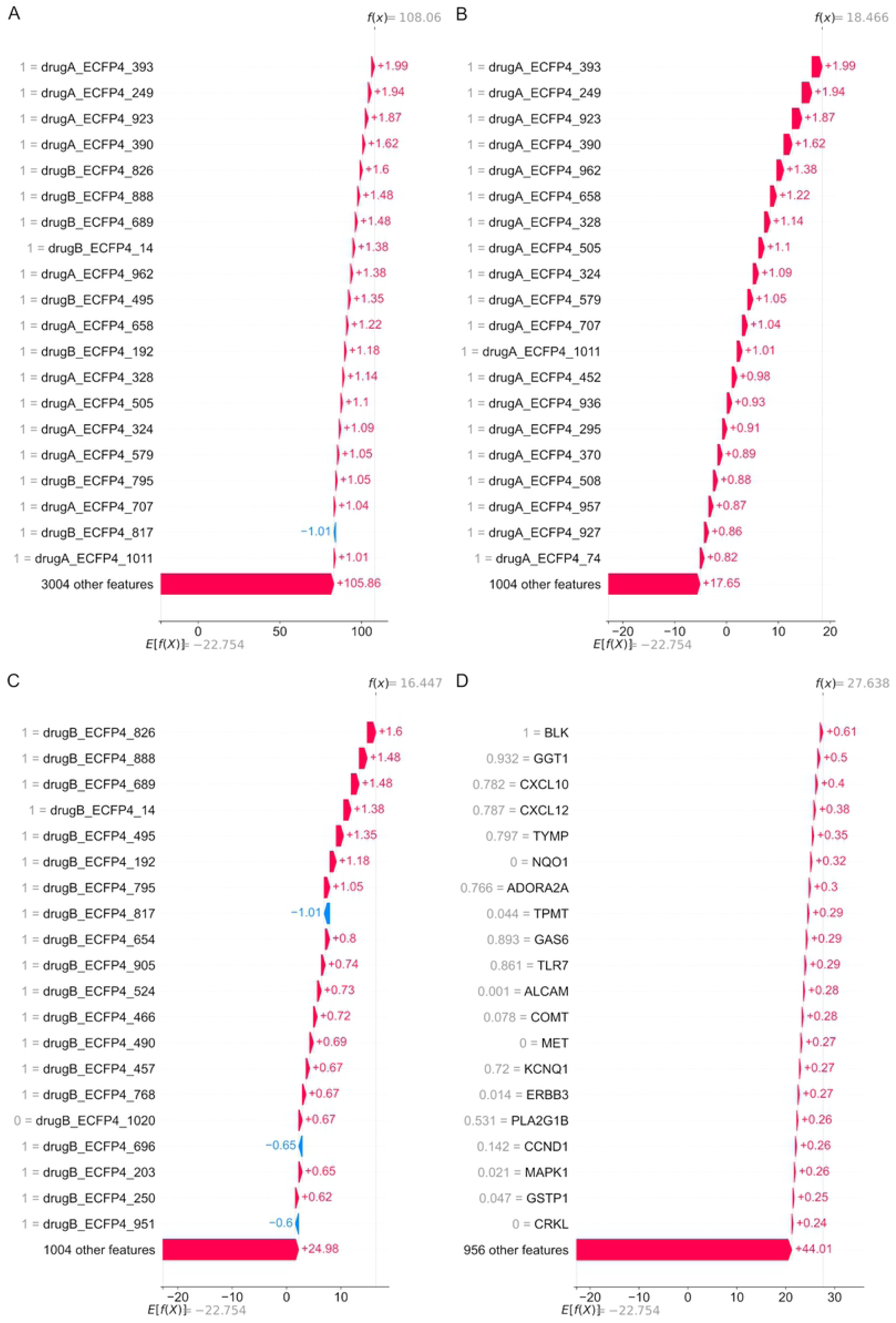
Most important features (ranked by absolute SHAP values) for test set example 11835, shown as waterfall plots. The plot starts at a base value of -22.754, which is the expected model output (determined based on a background dataset). Each row shows how each feature positively or negatively contributes to move the value from the expected output to the predicted value for this sample. (A) Top 20 features (from all features). (B) Top 20 drugA features. (C) Top 20 drugB features. (D) Top 20 gene expression features.

The SHAP values for each of the different input types were also analyzed separately. Fig 4B and Fig 4C show the 20 most important ECFP4 bits for drugA (Topotecan hydrochloride) and drugB (Gefitinib), respectively, along with the corresponding SHAP values. The 2D structures of 15 of the most important “ON” bits (value=1) that contributed positively are shown in S2 Fig for Topotecan and S3 Fig for Gefitinib.

As shown in S2 Fig, bits 249 and 658 capture parts of the lactone ring in Topotecan which establishes an important interaction with Topoisomerase I via hydrogen bonding [50]. Many of the remaining bits capture parts of the planar ring system that allows the drug to intercalate between two DNA base pairs and establish pi stacking interactions with the neighboring DNA bases [50]. These results reveal that our model was able to correctly identify which substructures are most important for the bioactivity of Topotecan.

The DL model was also able to identify features in Gefitinib that are more predictive of drug response and are consistent with the literature (S3 Fig). Bits 795, 490 and 203 capture parts of the quinazoline core that are involved in an important drug-protein interaction, with bit 203 focusing on the nitrogen atom that interacts with the adenosine triphosphate (ATP) binding site of EGFR via a hydrogen bond [52]. The majority of the bits in S3 Fig, however, correspond to the morpholine substituent or the 3-chloro-4-fluoro aniline substituent, which are not directly involved in drug-protein interactions [52]. The DL model may have identified these features as more important because they might be structures that distinguish Gefitinib from other compounds in the ALMANAC dataset, instead of being important for the drug activity *per se*.

The top 20 gene expression features are shown in Fig 4D. None of the genes that encode the primary targets of Topotecan and Gefitinib are found among the top 20 gene expression features. *TOP1*, which encodes DNA Topoisomerase I, was not present in the list of drug-gene interactions for Topotecan hydrochloride in DGIdb. The gene encoding DNA Topoisomerase I Mitochondrial (*TOP1MT*) was present in the list of drug-gene interactions, but it was only the 246th most important gene expression feature. EGFR ranked higher, being the 46th most important gene expression feature in this experiment. Additionally, none of the genes involved in the drug-gene interactions with Topotecan hydrochloride listed in DGIdb are among the top 20 most important gene expression features. Four of the top 20 genes (*BLK, MET, ERBB3* and *CRKL*) are involved in drug-gene interactions with Gefitinib, however. Gefitinib has been experimentally shown to be capable of binding to *BLK* [53], although with a much lower affinity than for EGFR. Since the expression of *BLK* is high in the SR cell line, the high SHAP value of *BLK* might be an indication that this interaction may be important for the response of the cells to Gefitinib. Amplification of *MET* [54] or *CRKL* [55] has been linked to acquired resistance to EGFR inhibitors, and *ERBB3* has also been implicated in resistance to EGFR tyrosine kinase inhibitors [54]. Since these genes are less expressed in the SR cell line, the higher SHAP values may be an indication that lower expression of these genes is essential for the cell line to be responsive to treatment with Gefitinib.

Interestingly, several of the most important genes have some kind of connection to the immune system. *CXCL10* and *CXCL12* are both chemokines involved in the recruitment of lymphocytes [56, 57]. *ADORA2A* plays a role in the regulation of the immune system [58], *TLR7* is involved in immune response [59], and *ALCAM* also plays a role in immunity [60]. It is possible that the DL identified these features as being essential for the prediction of drug synergy because the expression of these genes uniquely identifies the cell line in some way, and not necessarily due to the relevance of these genes for drug response.

A pre-ranked gene set enrichment analysis (GSEA) [61] was performed for test set example 11835 to determine if there is an enrichment of specific gene ontology (GO) terms among the most important genes. Genes were ranked based on their SHAP values. In total, the analysis identified 28 enriched gene sets, most of which refer to cell motility, cell death, and the regulation of kinase or transferase activity (S3 Table). We found that the gene sets with the highest normalized enrichment score (NES) values are leukocyte/lymphocyte-specific GO terms (S3 Table). We also observed that several of the enriched gene sets include the *EGFR* gene, which encodes the protein targeted by Gefitinib (S3 Table).

## Discussion

The results of this study suggest that drug features are more predictive of drug combination effects than cell line features, at least for the ALMANAC dataset, in line with previous results [19]. Inclusion of other cell line features besides gene expression data was not beneficial. Similar results were obtained in other drug response prediction studies [62, 63].

Several of the compound representation methods tested in this work produced models with similar performance. This finding is in agreement with a recent study by our group that compared the performance of different compound representations across different drug response prediction tasks and concluded that most representations perform similarly [68]. Since different drug representations may capture different characteristics of the compounds that may be equally predictive of drug response, the combined use of different types of drug representations might lead to an improvement in model performance and could be an interesting strategy to explore in the future.

Using prior biological knowledge for feature selection proved to be beneficial. Besides the cancer-specific and DGI gene lists that were utilized in this study, other cancer-specific gene lists and combinations of lists might provide even better results. Pathway propagation methods have been employed in other drug response prediction studies to simulate response to treatment [69] and to extend the selection of genes beyond known drug targets [26]. This strategy was not explored in the current study, but it might also be a way to improve the predictive capacity of the models. It is important to note that using drug targets or DGIs to select genes may limit the generalizability of the model, as new compounds may have DGIs that are different from the ones that were considered relevant for the training set.

Another approach that was not explored in this study is directly including biological knowledge in the neural network by representing the experiments as heterogeneous graphs and modeling drug response using GNNs. Since the number of unique compounds and cell lines in the ALMANAC dataset is relatively small, it is unclear if the models would benefit from the added complexity of this approach.

Despite training the models on a relatively large screening dataset, comprising over 200,000 screening experiments, there are only 59 cell lines and 104 drugs in ALMANAC. The number of unique compounds screened in the ALMANAC project is relatively small, a fact that might explain why fingerprint-based models performed slightly better than learned representations in this case, as learned representations are known to struggle when faced with smaller compound sets [64]. Considering the small number of unique cell lines and drugs, pre-training the feature encoding subnetworks using larger sets of compounds and larger omics datasets obtained from other sources could help improve model performance. Training models on larger and more diverse drug combination datasets from databases such as DrugCombDB [65] or DrugComb [66], which integrate data from several high-throughput screening experiments, could be another strategy to obtain models with better generalization capacity. However, it is important to note that different screening protocols might make it difficult to compare the synergy scores calculated for different screening datasets and to use them jointly to train models. Furthermore, integrating omics datasets from different studies requires the selection of adequate batch correction methods, which might not be straightforward [67].

We found that creating simple heterogeneous ensembles comprising both DL and ML models can improve performance. Preto et al. had previously observed that heterogeneous ensembles produced better classifiers than individual models when using the ALMANAC dataset [24], and other synergy prediction studies using different datasets also found that combining multiple models is beneficial [8, 10].

Although SHAP has allowed us to gain more insight into the model and to try to interpret its predictions in a biologically meaningful way, it is still difficult to explain how all of the most important features interact in a particular screening experiment and give rise to drug synergy. Interpretability methods allow us to explain how a DL model works, but they do not necessarily explain the underlying biology/chemistry. It is, therefore, unlikely that the methodology used in our study will be able to uncover the mechanisms underlying drug synergy for particular *< cell line− drugA− drugB >* triplets, but our findings do confirm that DL models are able to learn to associate important structural features of the compounds and tissue-specific gene expression patterns with drug response.

One important aspect to be considered is the validation scheme that is chosen. In our study, the data were randomly split into training, validation and test sets. Using a different validation scheme, such as a leave-drug-combinations-out approach (i.e. drug combinations that appear in one set are not included in the other sets), or even a leave-drugs-out or leave-cell-lines out validation scheme would likely result in lower performance scores. In addition, the models were validated with data from within the same drug combination screening study. Although this is a common model evaluation strategy, cross-validation within a single study has been shown to overestimate model generalizability [73]. In the future it would be interesting to assess different DL modeling strategies by performing a cross-study evaluation of the models as well [73].

## Materials and methods

### Datasets and data preprocessing

ALMANAC drug response data in the form of ComboScores for *< cell line, drugA, drugB >* triplets were downloaded from CellMiner Cross Database (CellMinerCDB) [74] (version 1.2).

Since ComboScore predictions should be independent of the order of the drugs in a given combination, we considered reverse drug order *< cell line, drugB, drugA >* triplets to be duplicates and the ComboScores for experiments involving the same cell line and same drug combination were averaged. The MDA-MB-468 cell line that is present in the original ALMANAC dataset was not considered in our analysis as it does not appear in the omics datasets we used in this study. The MDA-N cell line present in the omics datasets from CellMinerCDB was removed as it was not included in the ALMANAC project.

National Service Center (NSC) compound identifiers were used to map compounds to their respective SMILES strings using a compound structure-data file (SDF) file provided by NCI Developmental Therapeutics Program (DTP) for the ALMANAC dataset. For the compounds that were not successfully mapped using the previously described method, PubChemPy (version 1.0.4) was used to obtain canonical SMILES strings from PubChem [75] based on drug names. The SMILES strings were then preprocessed using the ChEMBL Structure Pipeline (version 1.0.0) [76], which removed salts and standardized the molecules according to ChEMBL-defined rules. The preprocessed SMILES were then used to compute different compound representations, depending on the type of architecture that was chosen for the drug subnetwork.

RNA-Seq gene expression data (log2(FPKM+1)) for the NCI-60 cell lines were also downloaded from CellMinerCDB [74]. Genes with constant expression values across all cell lines were removed. The expression dataset was then filtered using specific gene lists, as detailed in the Results section.

Besides the use of specific gene sets to reduce the number of features in the gene expression dataset, several other dimensionality reduction methods were also tested. UMAP was used to further reduce the dimensionality of the gene expression dataset filtered by protein coding genes (*expr*_*protein coding*_ dataset). Embeddings with 50 components were generated using the UMAP Python package (version 0.5.1) [40]. The algorithm was fit on the training data and then used to transform the training, validation and test sets. The datasets were min-max scaled beforehand. A total of 20 neighbors were considered when learning the manifold structure of the data. Pearson correlation was used as the distance metric, and the minimum distance between two points in the embedding was kept at the default value of 0.1.

We also evaluated the use of WGCNA [41] as a dimensionality reduction technique for gene expression data. The WGCNA R package (version 1.69) was used to find a total of 136 gene co-expression modules. Module eigengenes, which summarize the expression of genes in each module, were then used as input features instead of the original gene expression values.

When preparing the gene expression (*expr*_*protein coding*_) data to be fed into 1D or 2D CNN, two different methods were used to rearrange the genes so that their locations within the input tensors might be more biologically meaningful. The first method sorted the genes according to their chromosome positions, an approach similar to the method described in [77]. The second method employed hierarchical clustering to find a new order for the genes. The clustering algorithm used Pearson correlation as the distance metric and complete linkage. A dendrogram was created based on the clustering results, and the positions of the leaves (genes) was used to reorder the gene expression features. The gene expression data were then reshaped into the required formats for 1D and 2D CNNs.

A mutation annotation format (MAF) file containing mutation data from whole-exome sequencing for NCI-60 cell lines was downloaded from CBioPortal [78, 79]. Silent mutations and mutations in non-coding regions were excluded. The remaining data were then binarized and summarized at the gene level, with ‘1’ indicating that a given cell line had at least one alteration in a particular gene and zero indicating the absence of mutations. To create the gene-level mutation dataset, the data were further filtered using a list of drug-gene interactions from DGIdb, leaving a total of 636 genes. To create the pathway-level dataset, a new binary event matrix was created, where a ‘1’ indicates that a given cell line had at least one alteration in a given pathway. Genes were mapped to pathways using Molecular Signatures Database (MSigDB) [80, 81] (version 7.2) gene sets derived from the Reactome [82] database, which have been filtered to reduce the redundancy between gene sets. The pathway-level mutation dataset had a total of 1,510 features.

Genomic Identification of Significant Targets in Cancer (GISTIC)2 putative CNVs for the NCI-60 cell lines were also obtained from CBioPortal [78, 79]. Only genes that were present in the drug-gene interactions list obtained from DGIdb were kept, leaving a total of 952 features.

The omics datasets were individually merged with the drug response dataset to create the final omics datasets containing the corresponding cell line data for each experiment. Afterwards, the data were split into three subsets - a training set comprising around 80% of the data (239,943 examples), and validation and test sets each corresponding to approximately 10% of the original data (30,007 and 30,001 examples, respectively). After splitting the expression dataset into training/validation/test sets, the features were scaled to a range between zero and one (min-max scaling), with the exception of the UMAP and WGCNA-transformed data, which were not scaled after the dimensionality reduction step.

### Models

The DL models were built using a multimodal approach, in which each data type has its own corresponding feature-encoding subnetwork. The learned representations from each subnetwork were then concatenated into a single tensor before being fed into a final fully connected drug synergy prediction subnetwork.

Several different architectures were tested for each of the subnetworks. The gene expression subnetworks were either fully connected networks, 1D CNNs, or 2D CNNs. When included, the mutation and CNV subnetworks were always fully connected. For the drug subnetworks, we tested fully connected networks (when using ECFP4 fingerprints, LayeredFP or MTEs as inputs), GCN [45] subnetworks, GAT [46] subnetworks, and TextCNN [43] subnetworks (as implemented in DeepChem [44]). Each drug in a combination had a separate feature-encoding subnetwork, with weights being shared between the two drug subnetworks. The prediction subnetwork was always entirely composed of fully connected layers, ending in a single output unit with a linear activation function, given that the prediction task was approached as a regression problem.

All of the models were implemented using the Keras subpackage in Tensorflow [83] (version 2.2.0). The GCN and GAT subnetworks were built using graph layers implemented in the Spektral package [84] (version 1.0.7).

### Model hyperparameters and hyperparameter optimization

All models used the mean squared error (MSE) as the loss function, and the adaptive moment estimation (Adam) algorithm as the optimization method. Models were trained for a maximum of 500 epochs, using the EarlyStopping callback with a patience of 15 epochs to stop the learning process once the validation loss stopped improving. The mini-batch size was 64.

A selection of model hyperparameters, including the learning rate, the hidden layer activation function, the dropout rate and the L2 regularization penalty, as well as subnetwork-specific hyperparameters, were optimized for each combination of input data types and subnetwork architectures that we evaluated in this work. Hyperparameters were tuned using the validation set and the Bayesian Optimization with HyperBand (BOHB) algorithm [85] as implemented in Ray Tune (version 1.0.1). The hyperparameter search space was explored using Bayesian optimization [86] and the HyperBand [87] algorithm was used to stop underperforming trials early. A total of 50 configurations were evaluated for each model. The set of hyperparameters that minimized the validation MSE was considered the best configuration. More details on the hyperparameter search grids that were explored and the hyperparameter values that were chosen for each model are provided in S1 File.

### Model evaluation

After tuning the model hyperparameters, the models were evaluated on the independent test set. Model performance was evaluated using several scoring metrics, including the R^2^ and Spearman’s rank correlation coefficient.

In addition to comparing DL models with different omics and drug subnetworks, our models were also compared against non-DL models. Elastic net linear regression [88], SVR [89], and RF [90] models were implemented using the scikit-learn Python package (version 0.22.1) [91]. Since kernelized SVR does not scale well to larger datasets, we used the Nystroem method [92] to approximate the feature mappings of a RBF kernel before training a linear SVR on these approximations. An XGBoost [93] model was implemented using the xgboost Python package (version 1.4.0), and a LGBM [94] model was implemented using the lightgbm package (version 3.2.1). The ML models were trained on ECFP4 fingerprints and the gene expression values of DGI genes based on information from DGIdb. The input features were concatenated into a single dataset before training. Hyperparameters were optimized using Bayesian optimization.

The models were also compared against a baseline model that always predicts the average ComboScore value of the training set, and against models where the cell line and/or drug features were substituted with the corresponding one-hot encoded identifiers (*cell line*_*one hot*_ *+ drugs*_*one hot*_, *cell line*_*one hot*_ *+ drugs*_*ECFP4*_, and *expr*_*DGI*_ *+ drugs*_*one hot*_).

### Feature importance

The SHAP Python package (version 0.39.0) [48] was used to explain predictions made by the best DL models developed in this work. More specifically, we used Deep SHAP, which uses the Deep Learning Important FeaTures (DeepLIFT) [95] additive feature attribution method to approximate SHAP values. Feature importance was then determined based on these values. The most important fingerprint bits were visualized using drawing functions available in RDKit (version 2020.09.1.0).

A pre-ranked GSEA [61] was performed for test set example 11835 using the clusterProfiler R package (version 3.18.1) [96]. SHAP values were used to rank the genes prior to the analysis.

## Supporting information

**S1 Fig. The different types of deep learning and machine learning models developed in this study**.

**S1 Table. Performance scores for all of the models developed in this study**.

**S2 Table. Top 20 features (ranked by Gini importance) for the random forest model**.

**S2 Fig. 2D depiction of the top 15 most important ECFP4 “ON” bits with a positive effect on the model output for drugA (Topotecan) in test set example 11835**.

**S3 Fig. 2D depiction of the top 15 most important ECFP4 “ON” bits with a positive effect on the model output for drugB (Gefitinib) in test set example 11835**.

**S3 Table. Pre-ranked GSEA results**.

**S1 File. Hyperparameter search grids and selected hyperparameter values for each model**.

## Acknowledgments

This study was supported by the Portuguese Foundation for Science and Technology (FCT) under the scope of the strategic funding of UIDB/04469/2020 unit and through a PhD scholarship (SFRH/BD/130913/2017) awarded to Delora Baptista.

## References

1. Holohan C, Van Schaeybroeck S, Longley DB, Johnston PG. Cancer drug resistance: an evolving paradigm. Nature Reviews Cancer. 2013;13(10):714–726. doi:10.1038/nrc3599.

2. Dagogo-Jack I, Shaw AT. Tumour heterogeneity and resistance to cancer therapies. Nature Reviews Clinical Oncology. 2018;15(2):81–94. doi:10.1038/nrclinonc.2017.166.

3. Chatterjee N, Bivona TG. Polytherapy and Targeted Cancer Drug Resistance. Trends in Cancer. 2019;5(3):170–182. doi:10.1016/j.trecan.2019.02.003.

4. Al-Lazikani B, Banerji U, Workman P. Combinatorial drug therapy for cancer in the post-genomic era. Nature Biotechnology. 2012;30(7):679–692. doi:10.1038/nbt.2284.

5. Tallarida RJ. Quantitative Methods for Assessing Drug Synergism. Genes & Cancer. 2011;2(11):1003–1008. doi:10.1177/1947601912440575.

6. O’Neil J, Benita Y, Feldman I, Chenard M, Roberts B, Liu Y, et al. An Unbiased Oncology Compound Screen to Identify Novel Combination Strategies. Molecular Cancer Therapeutics. 2016;15(6):1155–1162. doi:10.1158/1535-7163.MCT-15-0843.

7. Holbeck SL, Camalier R, Crowell JA, Govindharajulu JP, Hollingshead M, Anderson LW, et al. The National Cancer Institute ALMANAC: A Comprehensive Screening Resource for the Detection of Anticancer Drug Pairs with Enhanced Therapeutic Activity. Cancer Research. 2017;77(13):3564–3576. doi:10.1158/0008-5472.CAN-17-0489.

8. Menden MP, Wang D, Mason MJ, Szalai B, Bulusu KC, Guan Y, et al. Community assessment to advance computational prediction of cancer drug combinations in a pharmacogenomic screen. Nature Communications. 2019;10(1):2674. doi:10.1038/s41467-019-09799-2.

9. Shoemaker RH. The NCI60 human tumour cell line anticancer drug screen. Nature Rev. 2006;6(10):813–823. doi:10.1038/nrc1951.

10. Bansal M, Yang J, Karan C, Menden MP, Costello JC, Tang H, et al. A community computational challenge to predict the activity of pairs of compounds. Nature Biotechnology. 2014;32(12):1213–1222. doi:10.1038/nbt.3052.

11. Gayvert KM, Aly O, Platt J, Bosenberg MW, Stern DF, Elemento O. A Computational Approach for Identifying Synergistic Drug Combinations. PLOS Computational Biology. 2017;13(1):e1005308. doi:10.1371/journal.pcbi.1005308.

12. Celebi R, Bear Don’t Walk O, Movva R, Alpsoy S, Dumontier M. In-silico Prediction of Synergistic Anti-Cancer Drug Combinations Using Multi-omics Data. Scientific Reports. 2019;9(1):8949. doi:10.1038/s41598-019-45236-6.

13. Sidorov P, Naulaerts S, Ariey-Bonnet J, Pasquier E, Ballester PJ. Predicting Synergism of Cancer Drug Combinations Using NCI-ALMANAC Data. Frontiers in Chemistry. 2019;7. doi:10.3389/fchem.2019.00509.

14. Nakano T, Jb B. Prediction of Compound Cytotoxicity Based on Compound Structures and Cell Line Molecular Characteristics. Journal of Computer Aided Chemistry. 2020;21:1–10.

15. Zagidullin B, Wang Z, Guan Y, Pitkänen E, Tang J. Comparative analysis of molecular fingerprints in prediction of drug combination effects. Briefings in Bioinformatics. 2021;doi:10.1093/bib/bbab291.

16. LeCun Y, Bengio Y, Hinton G. Deep learning. Nature. 2015;521(7553):436–444. doi:10.1038/nature14539.

17. Bengio Y, Courville A, Vincent P. Representation Learning: A Review and New Perspectives. IEEE Transactions on Pattern Analysis and Machine Intelligence. 2013;35(8):1798–1828. doi:10.1109/TPAMI.2013.50.

18. Preuer K, Lewis RPI, Hochreiter S, Bender A, Bulusu KC, Klambauer G. DeepSynergy: predicting anti-cancer drug synergy with Deep Learning. Bioinformatics. 2018;34(9):1538–1546. doi:10.1093/bioinformatics/btx806.

19. Xia F, Shukla M, Brettin T, Garcia-Cardona C, Cohn J, Allen JE, et al. Predicting tumor cell line response to drug pairs with deep learning. BMC Bioinformatics. 2018;19(S18):486. doi:10.1186/s12859-018-2509-3.

20. Zhang T, Zhang L, Payne PRO, Li F. In: Markowitz J, editor. Synergistic Drug Combination Prediction by Integrating Multiomics Data in Deep Learning Models. New York, NY: Springer US; 2021. p. 223–238.

21. Kuru HI, Tastan O, Cicek E. MatchMaker: A Deep Learning Framework for Drug Synergy Prediction. IEEE/ACM Transactions on Computational Biology and Bioinformatics. 2021; p. 1–1. doi:10.1109/TCBB.2021.3086702.

22. Kim Y, Zheng S, Tang J, Jim Zheng W, Li Z, Jiang X. Anticancer drug synergy prediction in understudied tissues using transfer learning. Journal of the American Medical Informatics Association. 2021;28(1):42–51. doi:10.1093/jamia/ocaa212.

23. Wang J, Liu X, Shen S, Deng L, Liu H. DeepDDS: deep graph neural network with attention mechanism to predict synergistic drug combinations. Briefings in Bioinformatics. 2021;doi:10.1093/bib/bbab390.

24. Preto AJ, Matos-Filipe P, Mourão J, Moreira IS. SynPred: Prediction of Drug Combination Effects in Cancer using Full-Agreement Synergy Metrics and Deep Learning. Preprints. 2021;.

25. Zhang H, Feng J, Zeng A, Payne P, Li F. Predicting tumor cell response to synergistic drug combinations using a novel simplified deep learning model. In: AMIA Annual Symposium Proceedings. vol. 2020. American Medical Informatics Association; 2020. p. 1364.

26. Liu Q, Xie L. TranSynergy: Mechanism-driven interpretable deep neural network for the synergistic prediction and pathway deconvolution of drug combinations. PLOS Computational Biology. 2021;17(2):e1008653. doi:10.1371/journal.pcbi.1008653.

27. Vaswani A, Shazeer N, Parmar N, Uszkoreit J, Jones L, Gomez AN, et al. Attention is all you need. In: Advances in neural information processing systems; 2017. p. 5998–6008.

28. Bazgir O, Ghosh S, Pal R. Investigation of REFINED CNN ensemble learning for anti-cancer drug sensitivity prediction. Bioinformatics. 2021;37(Supplement 1):i42.–i50. doi:10.1093/bioinformatics/btab336.

29. Jiang P, Huang S, Fu Z, Sun Z, Lakowski TM, Hu P. Deep graph embedding for prioritizing synergistic anticancer drug combinations. Computational and Structural Biotechnology Journal. 2020;18:427–438. doi:10.1016/j.csbj.2020.02.006.

30. Dong Z, Zhang H, Chen Y, Li F. Interpretable Drug Synergy Prediction with Graph Neural Networks for Human-AI Collaboration in Healthcare; 2021. Available from: http://arxiv.org/abs/2105.07082.

31. Yang J, Xu Z, Wu WKK, Chu Q, Zhang Q. GraphSynergy: a network-inspired deep learning model for anticancer drug combination prediction. Journal of the American Medical Informatics Association. 2021;28(11):2336–2345. doi:10.1093/jamia/ocab162.

32. Rozemberczki B, Gogleva A, Nilsson S, Edwards G, Nikolov A, Papa E. MOOMIN: Deep Molecular Omics Network for Anti-Cancer Drug Combination Therapy; 2021. Available from: http://arxiv.org/abs/2110.15087.

33. Ma J, Motsinger-Reif A. Prediction of synergistic drug combinations using PCA-initialized deep learning. BioData Mining. 2021;14(1):46. doi:10.1186/s13040-021-00278-3.

34. Bliss CI. The Toxicity of Poisons Applied Jointly. Annals of Applied Biology. 1939;26(3):585–615. doi:10.1111/j.1744-7348.1939.tb06990.x.

35. Subramanian A, Narayan R, Corsello SM, Peck DD, Natoli TE, Lu X, et al. A Next Generation Connectivity Map: L1000 Platform and the First 1,000,000 Profiles. Cell. 2017;171(6):1437–1452.e17. doi:10.1016/j.cell.2017.10.049.

36. Freshour SL, Kiwala S, Cotto KC, Coffman AC, McMichael JF, Song JJ, et al. Integration of the Drug–Gene Interaction Database (DGIdb 4.0) with open crowdsource efforts. Nucleic Acids Research. 2021;49(D1):D1144–D1151. doi:10.1093/nar/gkaa1084.

37. Tate JG, Bamford S, Jubb HC, Sondka Z, Beare DM, Bindal N, et al. COSMIC: the Catalogue Of Somatic Mutations In Cancer. Nucleic Acids Research. 2019;47(D1):D941–D947. doi:10.1093/nar/gky1015.

38. Sondka Z, Bamford S, Cole CG, Ward SA, Dunham I, Forbes SA. The COSMIC Cancer Gene Census: describing genetic dysfunction across all human cancers. Nature Reviews Cancer. 2018;18(11):696–705. doi:10.1038/s41568-018-0060-1.

39. Repana D, Nulsen J, Dressler L, Bortolomeazzi M, Venkata SK, Tourna A, et al. The Network of Cancer Genes (NCG): a comprehensive catalogue of known and candidate cancer genes from cancer sequencing screens. Genome Biology. 2019;20(1):1. doi:10.1186/s13059-018-1612-0.

40. McInnes L, Healy J, Melville J. UMAP: Uniform Manifold Approximation and Projection for Dimension Reduction; 2018. Available from: http://arxiv.org/abs/1802.03426.

41. Langfelder P, Horvath S. WGCNA: an R package for weighted correlation network analysis. BMC Bioinformatics. 2008;9(1):559. doi:10.1186/1471-2105-9-559.

42. Rogers D, Hahn M. Extended-Connectivity Fingerprints. Journal of Chemical Information and Modeling. 2010;50(5):742–754. doi:10.1021/ci100050t.

43. Kim Y. Convolutional Neural Networks for Sentence Classification. In: Proceedings of the 2014 Conference on Empirical Methods in Natural Language Processing (EMNLP). Stroudsburg, PA, USA: Association for Computational Linguistics; 2014. p. 1746–1751.

44. Ramsundar B, Eastman P, Walters P, Pande V, Leswing K, Wu Z. Deep Learning for the Life Sciences. O’Reilly Media; 2019.

45. Kipf TN, Welling M. Semi-Supervised Classification with Graph Convolutional Networks. In: 5th International Conference on Learning Representations, ICLR 2017, Toulon, France, April 24-26, 2017, Conference Track Proceedings. OpenReview.net; 2017. Available from: https://openreview.net/forum?id=SJU4ayYgl.

46. Velickovic P, Cucurull G, Casanova A, Romero A, Lio P, Bengio Y. Graph Attention Networks. In: 6th International Conference on Learning Representations, ICLR 2018, Vancouver, BC, Canada, April 30 - May 3, 2018, Conference Track Proceedings. OpenReview.net; 2018. Available from: https://openreview.net/forum?id=rJXMpikCZ.

47. Morris P, St Clair R, Hahn WE, Barenholtz E. Predicting Binding from Screening Assays with Transformer Network Embeddings. Journal of Chemical Information and Modeling. 2020;60(9):4191–4199. doi:10.1021/acs.jcim.9b01212.

48. Lundberg SM, Lee SI. A Unified Approach to Interpreting Model Predictions. In: Guyon I, Luxburg UV, Bengio S, Wallach H, Fergus R, Vishwanathan S, et al., editors. Advances in Neural Information Processing Systems 30. Curran Associates, Inc.; 2017. p. 4765–4774.

49. Ghandi M, Huang FW, Jané-Valbuena J, Kryukov GV, Lo CC, McDonald ER, et al. Next-generation characterization of the Cancer Cell Line Encyclopedia. Nature. 2019;569(7757):503–508. doi:10.1038/s41586-019-1186-3.

50. Staker BL, Hjerrild K, Feese MD, Behnke CA, Burgin AB, Stewart L. The mechanism of topoisomerase I poisoning by a camptothecin analog. Proceedings of the National Academy of Sciences. 2002;99(24):15387–15392.

51. Wakeling AE, Guy SP, Woodburn JR, Ashton SE, Curry BJ, Barker AJ, et al. ZD1839 (Iressa): an orally active inhibitor of epidermal growth factor signaling with potential for cancer therapy. Cancer research. 2002;62(20):5749–5754.

52. Yun CH, Boggon TJ, Li Y, Woo MS, Greulich H, Meyerson M, et al. Structures of Lung Cancer-Derived EGFR Mutants and Inhibitor Complexes: Mechanism of Activation and Insights into Differential Inhibitor Sensitivity. Cancer Cell. 2007;11(3):217–227. doi:10.1016/j.ccr.2006.12.017.

53. Davis MI, Hunt JP, Herrgard S, Ciceri P, Wodicka LM, Pallares G, et al. Comprehensive analysis of kinase inhibitor selectivity. Nature Biotechnology. 2011;29(11):1046–1051. doi:10.1038/nbt.1990.

54. Engelman JA, Zejnullahu K, Mitsudomi T, Song Y, Hyland C, Park JO, et al. MET Amplification Leads to Gefitinib Resistance in Lung Cancer by Activating ERBB3 Signaling. Science. 2007;316(5827):1039–1043. doi:10.1126/science.1141478.

55. Cheung HW, D. J, Boehm JS, He F, Weir BA, Wang X, et al. Amplification of CRKL Induces Transformation and Epidermal Growth Factor Receptor Inhibitor Resistance in Human Non–Small Cell Lung Cancers. Cancer Discovery. 2011;1(7):608–625. doi:10.1158/2159-8290.CD-11-0046.

56. Neville LF, Mathiak G, Bagasra O. The immunobiology of interferon-gamma inducible protein 10 kD (IP-10): A novel, pleiotropic member of the C-X-C chemokine superfamily. Cytokine & Growth Factor Reviews. 1997;8(3):207–219. doi:10.1016/S1359-6101(97)00015-4.

57. Bleul CC, Fuhlbrigge RC, Casasnovas JM, Aiuti A, Springer TA. A highly efficacious lymphocyte chemoattractant, stromal cell-derived factor 1 (SDF-1). Journal of Experimental Medicine. 1996;184(3):1101–1109. doi:10.1084/jem.184.3.1101.

58. Ohta A, Sitkovsky M. Extracellular Adenosine-Mediated Modulation of Regulatory T Cells. Frontiers in Immunology. 2014;5. doi:10.3389/fimmu.2014.00304.

59. Diebold SS, Kaisho T, Hemmi H, Akira S, Reis e Sousa C. Innate Antiviral Responses by Means of TLR7-Mediated Recognition of Single-Stranded RNA. Science. 2004;303(5663):1529–1531. doi:10.1126/science.1093616.

60. Bowen MA, Patel DD, Li X, Modrell B, Malacko AR, Wang WC, et al. Cloning, mapping, and characterization of activated leukocyte-cell adhesion molecule (ALCAM), a CD6 ligand. Journal of Experimental Medicine. 1995;181(6):2213–2220. doi:10.1084/jem.181.6.2213.

61. Subramanian A, Tamayo P, Mootha VK, Mukherjee S, Ebert BL, Gillette MA, et al. Gene set enrichment analysis: A knowledge-based approach for interpreting genome-wide expression profiles. Proceedings of the National Academy of Sciences. 2005;102(43):15545–15550. doi:10.1073/pnas.0506580102.

62. Costello JC, Heiser LM, Georgii E, Gönen M, Menden MP, Wang NJ, et al. A community effort to assess and improve drug sensitivity prediction algorithms. Nature Biotechnology. 2014;32(12):1202–1212. doi:10.1038/nbt.2877.

63. Iorio F, Knijnenburg TA, Vis DJ, Bignell GR, Menden MP, Schubert M, et al. A Landscape of Pharmacogenomic Interactions in Cancer. Cell. 2016;166(3):740–754. doi:10.1016/j.cell.2016.06.017.

64. Wu Z, Ramsundar B, Feinberg EN, Gomes J, Geniesse C, Pappu AS, et al. MoleculeNet: a benchmark for molecular machine learning. Chemical Science. 2018;9(2):513–530. doi:10.1039/C7SC02664A.

65. Liu P, Li H, Li S, Leung KS. Improving prediction of phenotypic drug response on cancer cell lines using deep convolutional network. BMC Bioinformatics. 2019;20(1):408. doi:10.1186/s12859-019-2910-6.

66. Zagidullin B, Aldahdooh J, Zheng S, Wang W, Wang Y, Saad J, et al. DrugComb: an integrative cancer drug combination data portal. Nucleic Acids Research. 2019;47(W1):W43–W51. doi:10.1093/nar/gkz337.

67. Goh WWB, Wang W, Wong L. Why Batch Effects Matter in Omics Data, and How to Avoid Them. Trends in Biotechnology. 2017;35(6):498–507. doi:10.1016/j.tibtech.2017.02.012.

68. Baptista D, Correia J, Pereira B, Rocha M. A Comparison of Different Compound Representations for Drug Sensitivity Prediction. In: Rocha M, Fdez-Riverola F, Mohamad Ms, Casado-Vara R, editors. Practical Applications of Computational Biology & Bioinformatics, 15th International Conference (PACBB 2021). Cham: Springer International Publishing; 2022. p. 145–154.

69. Li H, Li T, Quang D, Guan Y. Network Propagation Predicts Drug Synergy in Cancers. Cancer Research. 2018; p. canres.0740.2018. doi:10.1158/0008-5472.CAN-18-0740.

70. Elmarakeby HA, Hwang J, Arafeh R, Crowdis J, Gang S, Liu D, et al. Biologically informed deep neural network for prostate cancer discovery. Nature. 2021;598(7880):348–352. doi:10.1038/s41586-021-03922-4.

71. Hao J, Kim Y, Kim TK, Kang M. PASNet: pathway-associated sparse deep neural network for prognosis prediction from high-throughput data. BMC Bioinformatics. 2018;19(1):510. doi:10.1186/s12859-018-2500-z.

72. Gaudelet T, Malod-Dognin N, Sánchez-Valle J, Pancaldi V, Valencia A, Przulj N. Unveiling new disease, pathway, and gene associations via multi-scale neural network. PLOS ONE. 2020;15(4):e0231059. doi:10.1371/journal.pone.0231059.

73. Xia F, Allen J, Balaprakash P, Brettin T, Garcia-Cardona C, Clyde A, et al. A cross-study analysis of drug response prediction in cancer cell lines. Briefings in Bioinformatics. 2022;23(1). doi:10.1093/bib/bbab356.

74. Rajapakse VN, Luna A, Yamade M, Loman L, Varma S, Sunshine M, et al. CellMinerCDB for Integrative Cross-Database Genomics and Pharmacogenomics Analyses of Cancer Cell Lines. iScience. 2018;10:247–264. doi:10.1016/j.isci.2018.11.029.

75. Kim S, Chen J, Cheng T, Gindulyte A, He J, He S, et al. PubChem in 2021: new data content and improved web interfaces. Nucleic Acids Research. 2021;49(D1):D1388–D1395. doi:10.1093/nar/gkaa971.

76. Bento AP, Hersey A, Félix E, Landrum G, Gaulton A, Atkinson F, et al. An open source chemical structure curation pipeline using RDKit. Journal of Cheminformatics. 2020;12(1):51. doi:10.1186/s13321-020-00456-1.

77. Lyu B, Haque A. Deep Learning Based Tumor Type Classification Using Gene Expression Data. In: Proceedings of the 2018 ACM International Conference on Bioinformatics, Computational Biology, and Health Informatics. New York, NY, USA: ACM; 2018. p. 89–96.

78. Cerami E, Gao J, Dogrusoz U, Gross BE, Sumer SO, Aksoy BA, et al. The cBio Cancer Genomics Portal: An Open Platform for Exploring Multidimensional Cancer Genomics Data Cancer Discovery. 2012;2(5):401–404. doi:10.1158/2159-8290.CD-12-0095.

79. Gao J, Aksoy BA, Dogrusoz U, Dresdner G, Gross B, Sumer SO, et al. Integrative Analysis of Complex Cancer Genomics and Clinical Profiles Using the cBioPortal. Science Signaling. 2013;6(269):pl1.#x2013;pl1. doi:10.1126/scisignal.2004088.

80. Subramanian A, Tamayo P, Mootha VK, Mukherjee S, Ebert BL, Gillette MA, et al. Gene set enrichment analysis: A knowledge-based approach for interpreting genome-wide expression profiles. Proceedings of the National Academy of Sciences. 2005;102(43):15545–15550. doi:10.1073/pnas.0506580102.

81. Liberzon A, Subramanian A, Pinchback R, Thorvaldsdottir H, Tamayo P, Mesirov JP. Molecular signatures database (MSigDB) 3.0. Bioinformatics. 2011;27(12):1739–1740. doi:10.1093/bioinformatics/btr260.

82. Jassal B, Matthews L, Viteri G, Gong C, Lorente P, Fabregat A, et al. The reactome pathway knowledgebase. Nucleic Acids Research. 2019;doi:10.1093/nar/gkz1031.

83. Abadi M, Agarwal A, Barham P, Brevdo E, Chen Z, Citro C, et al. TensorFlow: Large-Scale Machine Learning on Heterogeneous Distributed Systems. Nature Neuroscience. 2016;16(4):486–492.

84. Grattarola D, Alippi C. Graph Neural Networks in TensorFlow and Keras with Spektral [Application Notes]. IEEE Computational Intelligence Magazine. 2021;16(1):99–106.

85. Falkner S, Klein A, Hutter F. BOHB: Robust and efficient hyperparameter optimization at scale. In: International Conference on Machine Learning. PMLR; 2018. p. 1437–1446.

86. Mo?ckus J. On Bayesian Methods for Seeking the Extremum. In: Optimization Techniques IFIP Technical Conference. Berlin, Heidelberg: Springer Berlin Heidelberg; 1975. p. 400–404.

87. Li L, Jamieson K, DeSalvo G, Rostamizadeh A, Talwalkar A. Hyperband: A novel bandit-based approach to hyperparameter optimization. The Journal of Machine Learning Research. 2017;18(1):6765–6816.

88. Zou H, Hastie T. Regularization and variable selection via the elastic net. Journal of the Royal Statistical Society: Series B (Statistical Methodology). 2005;67(2):301–320. doi:10.1111/j.1467-9868.2005.00503.x.

89. Drucker H, Burges CJC, Kaufman L, Smola AJ, Vapnik V. Support vector regression machines. In: Advances in neural information processing systems; 1997. p. 155–161.

90. Breiman L. Random forests. Machine Learning. 2001;45(1):5–32. doi:10.1023/A:1010933404324.

91. Pedregosa F, Varoquaux G, Gramfort A, Michel V, Thirion B, Grisel O, et al. Scikit-learn: Machine Learning in Python. Journal of Machine Learning Research. 2012;12(Oct):2825–2830.

92. Williams CKI, Seeger M. Using the Nyström Method to Speed up Kernel Machines. In: Proceedings of the 13th International Conference on Neural Information Processing Systems. NIPS’00. Cambridge, MA, USA: MIT Press; 2000. p. 661–667.

93. Chen T, Guestrin C. Xgboost: A scalable tree boosting system. In: Proceedings of the 22nd acm sigkdd international conference on knowledge discovery and data mining; 2016. p. 785–794.

94. Ke G, Meng Q, Finley T, Wang T, Chen W, Ma W, et al. Lightgbm: A highly efficient gradient boosting decision tree. Advances in neural information processing systems. 2017;30:3146–3154.

95. Shrikumar A, Greenside P, Kundaje A. Learning important features through propagating activation differences. In: International Conference on Machine Learning. PMLR; 2017. p. 3145–3153.

96. Yu G, Wang LG, Han Y, He QY. clusterProfiler: an R Package for Comparing Biological Themes Among Gene Clusters. OMICS: A Journal of Integrative Biology. 2012;16(5):284–287. doi:10.1089/omi.2011.0118.

